# Sleep spindles in primates: modelling the effects of distinct laminar thalamocortical connectivity in core, matrix, and reticular thalamic circuits

**DOI:** 10.1101/2022.04.27.489802

**Authors:** Arash Yazdanbakhsh, Helen Barbas, Basilis Zikopoulos

## Abstract

Sleep spindles are associated with the beginning of deep sleep and memory consolidation and are disrupted in schizophrenia and autism. In primates, distinct core and matrix thalamocortical (TC) circuits regulate sleep-spindle activity, through communications that are filtered by the inhibitory thalamic reticular nucleus (TRN) however, little is known about typical TC network interactions and the mechanisms that are disrupted in brain disorders. We developed a primate-specific, circuit-based TC computational model with distinct core and matrix loops that can simulate sleep spindles. We implemented novel multilevel cortical and thalamic mixing, and included local thalamic inhibitory interneurons, and direct layer 5 projections of variable density to TRN and thalamus to investigate the functional consequences of different ratios of core and matrix node connectivity contribution to spindle dynamics. Our simulations showed that spindle power in primates can be modulated based on the level of cortical feedback, thalamic inhibition, and engagement of model core vs. matrix, with the latter having a greater role in spindle dynamics. The study of the distinct spatial and temporal dynamics of core-, matrix-, and mix-generated sleep spindles establishes a framework to study disruption of TC circuit balance underlying deficits in sleep and attentional gating seen in autism and schizophrenia.

## Introduction

Sleep spindles are widespread oscillations associated with the beginning of deep NREM sleep, learning, and memory consolidation (Luthi, 2014; Steriade, 2005; Steriade et al., 1987). Because sleep spindles are disrupted in a variety of neurological and psychiatric conditions, they are an important clinical marker of atypical brain function in several disorders, including schizophrenia and autism (Farmer et al., 2018; Ferrarelli et al., 2007; Manoach et al., 2010; Mylonas et al., 2022).

Reciprocal connections between thalamic nuclei and cortical areas that are gated by the inhibitory thalamic reticular nucleus (TRN) form thalamocortical (TC) circuits that regulate sleep-spindle activity (Jones, 2007). However, in mammals two anatomically and functionally distinct TC circuits can be identified, the ‘core’ and the ‘matrix’ loops (Jones, 1998; Müller et al., 2020; Rovo et al., 2012; Zikopoulos and Barbas, 2007a), and it is not clear how spindle activity and associated functions are regulated by each circuit through synaptic, cellular, or regional interactions at the level of the cortex or thalamus. The core TC circuits, prevalent in sensory thalamus, drive activity focally in the middle cortical layers. In turn, these core thalamic neurons are innervated by small ‘modulatory’ cortical axon terminals from pyramidal neurons in layer 6 (L6). The matrix TC circuits, prevalent in high-order thalamus, have a complementary organization: a mix of large and small axon terminals from cortical layer 5 (L5) pyramidal neurons drive activity of matrix thalamic neurons that, in turn, innervate broadly and modulate the superficial cortical layers. The relative prevalence of the two TC loops differs, with core circuits prevailing in primary and unimodal association cortical areas and first-order relay nuclei, whereas matrix circuits predominate in association and limbic cortical areas and high-order thalamic nuclei (Harris and Shepherd, 2015; Jones, 1998; Rouiller and Welker, 2000; Rovo et al., 2012; Sherman and Guillery, 2006, 1996). However, there is considerable overlap of these parallel loops across TC networks, increasing the complexity of the system (Jones, 1998; Müller et al., 2020; Rovo et al., 2012; Zikopoulos and Barbas, 2007a).

Importantly, in addition to the excitatory connections in the thalamus, both TC circuits likely engage an extensive network of local thalamic inhibitory interneurons that have expanded significantly in evolution and constitute a hallmark of the primate thalamus (Arcelli et al., 1997), increasing the complexity of potential interactions in primates. Moreover, the connectivity of the inhibitory TRN with core and matrix TC loops is not well studied. The TRN is a key generator of spindle oscillations that intercepts all TC communications but sends inhibitory projections only back to the thalamus (Luthi, 2014; Steriade, 2005; Steriade et al., 1987). The TRN gets most of its cortical input from L6 pyramidal neurons that participate in core TC loops (Bourassa et al., 1995; Bourassa and Desche^nes, 1995; Kakei et al., 2001), but is predicted to get substantial input from L5 pyramidal neurons of matrix TC loops in primates (Zikopoulos and Barbas, 2007b, 2006), as was recently shown to some extent in mice (Hádinger et al., 2022; Hadinger et al., 2019; Prasad et al., 2020).

Based on this evidence, we set out to test the impact of local thalamic inhibition, variable levels of cortical L6 and L5 pyramidal neuron terminations in TRN, and the effects of these connections in core and matrix TC circuits and spindle activity, through the construction of a rate-based computational TC model. Our model successfully simulated relay and filtered signals to sustain and propagate spindle oscillations with different powers, depending on the level of cortical feedback, thalamic inhibition, and involvement of model core vs. matrix circuits. Our simulations additionally revealed differences in the spatiotemporal dynamics of core-generated, matrix-generated, or mixed types of spindles, highlighting novel circuit mechanisms involved in typical TC functions, like the sleep-wake cycle, sensory processing, attentional gating, and memory consolidation. Importantly, characterization of novel interactions between key nodes of TC circuits points to potential disruptions of mechanisms that may underlie atypical spindle dynamics in disorders, including autism, and schizophrenia.

## Methods

### Model design: basic connectivity frame

The model is based on key molecular, anatomical, and connectivity features of distinct TC circuits and their specialized interactions with the inhibitory TRN in primates that frame a basic TC circuit, as elaborated below, and summarized in Fig. 1.

**Figure 1.**
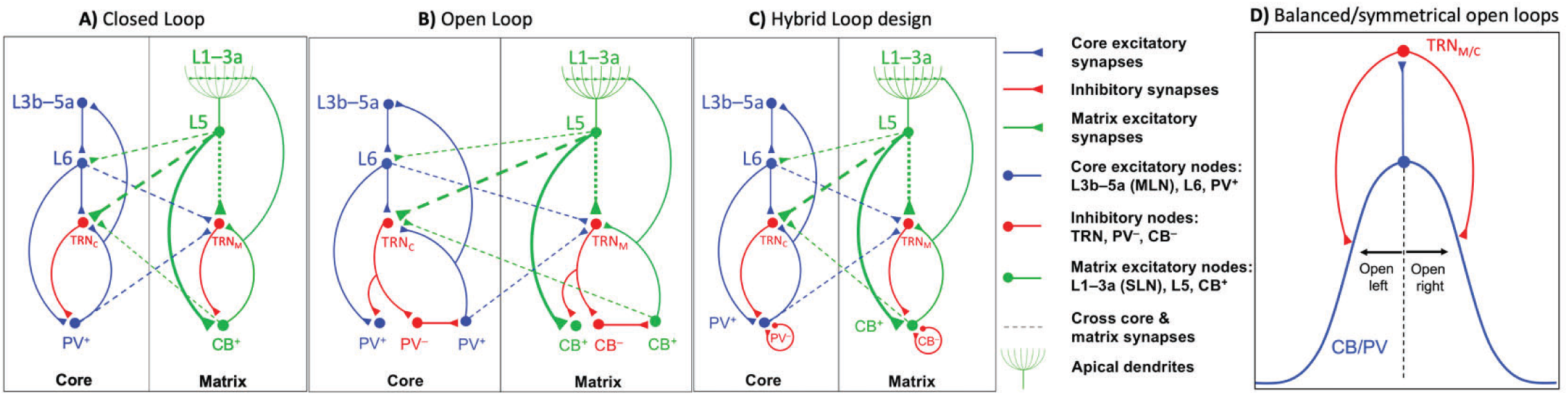
The proposed basic thalamocortical (TC) core, matrix and mix circuit. The model circuit was based on two parallel thalamocortical loops: Core - PV+ excitatory thalamic neurons project focally to the middle cortical layers and get feedback from cortical layer 6. Matrix - CB+ thalamic neurons innervate broadly the superficial cortical layers and receive projections from cortical layer 5. Layer 5 terminals in the thalamus are larger (thick arrowheads) than terminals from layer 6. Thalamic and cortical neurons project to TRN (TRN_C_ and TRN_M_). Mix - The two parallel loops cross-connect at the level of the thalamus and/or cortex. The mix dynamics were implemented in two different ways: 1) *corticoreticular*, where L6 and L5 concurrently excited TRN_M_ and TRN_C_, 2) *thalamoreticular*, where PV^+^ and CB^+^ concurrently excited TRN_M_ and TRN_C_, 3) cortical, where mixing of core and matrix networks occurred at the level of the cortex, through columnar laminar interactions, simplified as a cross-connection between L5 and L6 (dashed line); this way, we could modify core (L6 & PV^+^) vs. matrix (L5 & CB^+^) contribution in the mix. **(A)** In closed loop connections, TRN neurons directly inhibit the thalamic neurons that excite them. **B)** The anatomical circuits underlying open loop connections are shown, in which TRN neurons do not directly inhibit the thalamic neurons that excited them. **(C)** In the model we simulated the open loop architecture through implementation of the closed loop connections in which TRN neurons directly inhibit the thalamic neurons that excited them, following a symmetric Gaussian spread with the peak at the reciprocal thalamic neuron. Accordingly, the *bilateral* and symmetric Gaussian connectivity strength from TRN to thalamus resembles bilateral and symmetric open loops with opposite open directions (balanced). We called this the Hybrid Loop design. Inhibitory model neurons are shown in red, including TRN (TRN_C_ and TRN_M_ for core and matrix), PV-, and CB-local thalamic inhibitory interneurons. Blue units are core excitatory elements, i.e., neurons in cortical layers 4 and 6 (L3b-5a and L6), as well as thalamic PV^+^ neurons. Green units are matrix excitatory units, i.e., neurons in cortical layer 5 (L5), whose apical dendrites reach upper layers (L1-3a) and receive widespread input from excitatory thalamic CB^+^ neurons. **D)** When the Gaussian peak of connectivity strength is to the thalamic neuron which directly excites the TRN neuron, due to spatially symmetric bilateral Gaussian spread of connectivity strength, the circuit is composed of two open loops in opposite directions (balanced), then our architecture is functionally similar to a closed-loop.

There are two parallel functionally, structurally, and neurochemically distinct TC loops: the core and matrix (Jones, 2007). Thalamic neurons in core and matrix TC circuits of primates can be distinguished neurochemically (Clascá et al., 2012; Munkle et al., 2000). In core loops, excitatory TC projection neurons express the calcium-binding protein parvalbumin (PV). Core PV+ excitatory thalamic neurons project focally and drive activity of the middle cortical layers (mainly 4, but also 3b, and 5a) and get feedback from cortical layer 6. In the parallel matrix loops, excitatory TC projection neurons express the calcium-binding protein calbindin (CB). Matrix CB+ thalamic neurons innervate broadly and modulate primarily the superficial cortical layers (1-3a), recruiting a broad horizontal spread that targets apical dendrites of pyramidal cortical neurons from layers 2, 3, and 5, which extend to the upper cortical layers. Cortical layer 5 neurons target and drive activity of matrix CB+ thalamic neurons.

The entirely inhibitory TRN in primates covers almost the entire thalamus and gates all reciprocal connections between the cortex and the thalamus (Zikopoulos and Barbas, 2007b). The TRN receives input from excitatory thalamic projection neurons and pyramidal neurons in the cortex that project to the thalamus. Anatomical and physiological data suggest that cortical input drives TRN activity (Liu and Jones, 1999; Zhang and Jones, 2004). In turn, the TRN sends inhibitory projections only back to the thalamus. Until recently, it was thought that layer 6, but not layer 5 cortical pyramidal neurons project to TRN (Guillery, 2005, 1995; Guillery et al., 1998; Guillery and Harting, 2003; Guillery and Sherman, 2002; Kakei et al., 2001; Rouiller and Durif, 2004; Rouiller and Welker, 2000, 1991; Sherman and Guillery, 2002). We first provided strong indirect evidence for a projection from layer 5 to TRN, when we studied prefrontal corticothalamic projections and compared them with projections from sensory association cortices and other corticosubcortical projections in primates (Zikopoulos and Barbas, 2007a, 2006). Like other cortices, prefrontal areas project to the thalamus mainly from layer 6, but also issue significant projections from layer 5 (Xiao et al., 2009). We found that prefrontal layer 5 axon terminations in the thalamus constituted a mix of large and small boutons (thin and thick connections in Fig. 1) and overall, they were larger than terminals from sensory association areas known to originate from layer 6. Recent studies in mice confirmed that there are direct projections of variable density from L5 pyramidal neurons to TRN from some cortical areas, especially association areas that likely participate in matrix circuits (Hádinger et al., 2022; Hadinger et al., 2019; Prasad et al., 2020). Since the presence of L5 projections to TRN is not ubiquitous we represent it with a dotted line from L5 to TRN in the simplified diagram in Fig. 1, which shows the main nodes of core and matrix circuits.

There is evidence that TRN neurons are paired with thalamic neurons, forming reciprocal, closed-loop circuits, in which TRN neurons send GABAergic input to the thalamic neurons that innervate them directly (Fig. 1A) (Brown et al., 2020; FitzGibbon et al., 2000; Gentet and Ulrich, 2003; Hale et al., 1982; Lo and Murray Sherman, 1994; McAlonan et al., 2006; Pinault, 2004; Pinault and Deschenes, 1998; Sherman and Guillery, 1996; Shosaku, 1986; Steriade et al., 1993; Warren et al., 1994; Willis et al., 2015), reviewed in Edward G. Jones (Jones, 2007). In contrast, in open-loop circuits, TRN neurons innervate other thalamic neurons that do not directly innervate them (Fig. 1B) (Crabtree, 1998; Crabtree et al., 1998; Crabtree and Isaac, 2002; Kimura, 2014; Kimura et al., 2007; Lam and Murray Sherman, 2015; Lam and Sherman, 2005; Lee et al., 2010; McAlonan et al., 2006; Pinault and Deschenes, 1998).

In our model circuit we accounted for the TRN-thalamic loop architecture complexity, by representing innervation strength with Gaussian curves that have variable spread. We called this architecture Hybrid Loop Design (Fig. 1C). When the Gaussian peak of connectivity strength is to the thalamic neuron which directly excites the TRN neuron, due to spatially symmetric bilateral Gaussian spread of connectivity strength, the loop is composed of two open loops in opposite directions (balanced, Fig. 1D), then our architecture is functionally similar to a closed-loop. Conversely, when the connectivity spread is either spatially asymmetrical or follows a shifted Gaussian, then our circuit architecture resembles an open-loop. In addition, in primates, there is extensive presence of local inhibitory neurons in the thalamus, shown as PV- and CB-in Fig. 1 (Jones, 2007), which may participate in open-loop circuits.

The two parallel core and matrix circuits are not necessarily isolated but can overlap across TC networks. This overlap can be extensive in high-order TC networks, like the ones linking the mediodorsal nucleus (MD) or the ventral anterior nucleus (VA) with the prefrontal cortex (Jones, 1998; Müller et al., 2020; Rovo et al., 2012; Zikopoulos and Barbas, 2007a). However, it is not known whether this overlap is limited at the level of the thalamus, or if it is also present at the level of the cortex and TRN. To account for the potential overlap of core and matrix circuits at each TC node we included cross-connections (dashed lines in Fig. 1) that can facilitate *mixing* of the two parallel loops at all levels. For cortical core contribution in the mix, L6 directly projects to TRN_M_ and for cortical matrix contribution in the mixing, L5 directly projects to TRN_C_. For thalamic core contribution in the mix, PV^+^ directly projects to TRN_M_ and for thalamic matrix contribution in the mix, CB^+^ directly projects to TRN_C_. We also include mixing of core and matrix networks at the level of the cortex through several laminar interactions, simplified in Fig. 1 as a cross-connection between L5 and L6 (dashed line).

### Mathematical specification of the rate model: Dynamics of model neurons

We constructed a computational model of core and matrix TC circuits based on the connectivity above. We expressed each simulated neuron’s activity, based on inhibitory and excitatory inputs, over time, as a first order differential equation. Model neurons were represented by a single-compartment voltage (*v*) over time (Layton et al., 2014, 2012; Layton and Yazdanbakhsh, 2015) that obeys the following shunting equation:

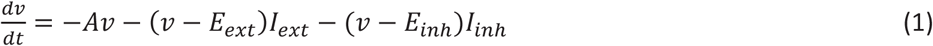

In this equation, *A* denotes the constant decay (leakage) rate, which brings *v* back to zero when there is no excitatory or inhibitory input to the neuron, to fulfil the physiological constraint that a neuron without input finally goes back to its resting potential. *I*_*ext*_ and *I*_*inh*_ specify the summed excitatory and inhibitory inputs to the neuron at each time. The terms *E*_*ext*_ and *I*_*inh*_ refer to excitatory, and inhibitory reversal potentials, respectively, which keep *v* within *E*_*ext*_ and *I*_*inh*_ range to fulfil the physiological constraint of the limited dynamical range of each neuron activity. We set *E*_*ext*_ and *I*_*inh*_ equal to 1 and -1; therefore, our model neurons activities fell within [-1, 1] range. Table 1 shows *I*_*ext*_ and *I*_*inh*_ for each model neuron type as illustrated in Fig. 1.

**Table 1.** Model neurons and their excitatory and inhibitory inputs (see Fig. 1)

| Model neuron | Symbol | Excitatory input ($I_{ext}$) | Inhibitory input ($I_{inh}$) |
| --- | --- | --- | --- |
| Matrix excitatory thalamic neurons | $CB_i^+$ | $\sum_{j=1}^N (L_5 \rightarrow CB^+)_{ji} L_5^j$ | $\sum_{j=1}^N (CB^- \rightarrow CB^+)_{ji} CB_{j-\eta}^- +$<br>$(TRN_M \rightarrow CB^+)_{ji} TRN_M^j$ |
| Core excitatory thalamic neurons | $PV_i^+$ | $\sum_{j=1}^N (L_6 \rightarrow PV^+)_{ji} L_6^j$ | $\sum_{j=1}^N (PV^- \rightarrow PV^+)_{ji} PV_{j-\eta}^- +$<br>$(TRN_C \rightarrow PV^+)_{ji} TRN_C^j$ |
| Matrix inhibitory thalamic neurons | $CB_i^-$ | $\sum_{j=1}^N (CB^+ \rightarrow CB^-)_{ji} CB_j^+$ | $\sum_{j=1}^N (TRN_M \rightarrow CB^-)_{ji} TRN_M^j$ |
| Matrix inhibitory thalamic neurons | $PV_i^-$ | $\sum_{j=1}^N (PV^+ \rightarrow PV^-)_{ji} PV_j^+$ | $\sum_{j=1}^N (TRN_C \rightarrow PV^-)_{ji} TRN_C^j$ |
| TRN Matrix | $TRN_M^i$ | $\sum_{j=1}^N (CB^+ \rightarrow TRN_M)_{ji} CB_{j+\eta}^+ + (L_5 \rightarrow$<br>$TRN_M)_{ji} L_5^j$ | $\sum_{j=1}^N (TRN_M \rightarrow TRN_M)_{ji} TRN_M^j$ |
| TRN Core | $TRN_C^i$ | $\sum_{j=1}^N (PV^+ \rightarrow TRN_C)_{ji} PV_{j+\eta}^+ + (L_6 \rightarrow$<br>$TRN_C)_{ji} L_6^j$ | $\sum_{j=1}^N (TRN_C \rightarrow TRN_C)_{ji} TRN_C^j$ |
| Superficial Layer<br>Neurons (SLN) | $L_{1-3a}^i$ | $\sum_{j=1}^N (CB^+ \rightarrow L_{1-3a})_{ji} CB_{j+\eta}^+$ | |
| Middle Layer<br>Neurons (MLN) | $L_{3b-4}^i$ | $\sum_{j=1}^N (PV^+ \rightarrow L_{3b-4})_{ji} PV_{j+\eta}^+$ | |
| Cortical Layer 5<br>neurons | $L_5^i$ | $\sum_{j=1}^N (L_{1-3a} \rightarrow L_5)_{ji} L_{1-3a}^j$ | |
| Cortical Layer 6<br>neurons | $L_6^i$ | $\sum_{j=1}^N (L_{3b-4} \rightarrow L_6)_{ji} L_{3b-4}^j$ | |

To facilitate description of excitatory/inhibitory interactions (*I*_*ext*_ and *I*_*inh*_) we constructed thalamic, reticular, and cortical networks as a distance-dependent on-center-off-surround shunting network, as we have described in the past (Layton et al., 2014, 2012; Layton and Yazdanbakhsh, 2015; Sherbakov and Yazdanbakhsh, 2013; Wurbs et al., 2013) and recently in computational models of cortico-thalamic-amygdalar circuits (John et al., 2016, 2013). Such distant dependent on-center interareal interactions are modeled by bell-shaped (Gaussian) profiles reflected in *I*_*ext*_ column of Table 1 by (*X → Y*) symbols, in which *X* and *Y* indicate the presynaptic and postsynaptic regions, for example, in row 1 of Table 1, *L*_*5*_ and *CB*^*+*^, respectively. Similarly, the off-surround distant dependent reflected in the *I*_*inh*_ column. Networks of this type offer a simple, biologically-plausible means to implement contrast-enhancement and a host of other processes, by varying the strength of the off-surround inhibition to modulate the tuning curve of each cell (Cao et al., 2015; Qian and Yazdanbakhsh, 2015). Strong inhibition yields sharp contrast, fine tuning, and high attention. Weak inhibition leads to spreading activity, lower contrast, and inattentiveness. These processes may be altered in autism and schizophrenia leading to deficits in attentional gating (Medalla and Barbas, 2009). Thus, modulating the balance between excitation and inhibition in various nodes of the circuit can provide a way to investigate the effects of various network elements on signal processing and attention.

### Equations for model neurons

Equation 1 is the general equation for all neurons of the model circuit of Fig. 1. The key distinction between model neurons is based on their excitatory (*I*_*ext*_) and inhibitory (*I*_*inh*_) inputs stemming from the neuroanatomy of the connectivity; therefore, the current neural model reflects the impact of neuroanatomy in generating distinct spatio-temporal dynamics of core and matrix TC loop spindles (Piantoni et al., 2016), based on the position and distribution of neurons rather than differences of reticular, thalamic, and cortical neuron types. Each row of Table 1 represents the presynaptic total excitatory and inhibitory input to each model neuron (i.e., CB^+^, TRN_M_, etc.). Indices *i* and *j* indicate the spatial position of model neurons *i* and *j* on a one-dimensional array. Pre- to post-synaptic connections are characterized by (Pre→Post) symbols sub-indexed by *ji* indicating the *j*th presynaptic to *i*th postsynaptic neuron strength. The relative strength of *j*th pre-synaptic to *i*th post-synaptic neuron follows a normal distribution centered at *i* (i.e., the *i*th pre- to *i*th post-synaptic connection is the strongest).

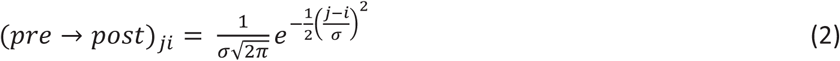

### Spindles in the model TRN

Neurons in the TRN send inhibitory GABAergic projections to thalamus (Fig. 1). TRN neurons can fire in bursts, generating IPSPs in excitatory thalamic neurons (PV+ and CB+), which in turn exhibit rebound bursts, activating the TRN neurons. By simulating neurons using the rate-based approach detailed above, we introduced burst-like activity input in our model TRN neurons (Fig. 2A), which mimics the temporal dynamic of constituent bursts in a spindle. The y-axis shows normalized values with the lower and upper limits of -1 and +1, similar to the lower and upper bounds of all of the model neurons’ activities. The inter-burst interval was set to 100ms to replicate a 10 Hz burst rate in 0.5 seconds. Such a voltage induction in model neurons is within the physiological range of sleep spindles with a frequency of 7-15 Hz and duration of 0.5 – 3 seconds (Luthi, 2014; Steriade, 2005; Steriade et al., 1987). In order to have the input temporal dynamics of Fig. 2A as close as possible to the burst sequence of spindles, we approximated Fig. 2B of (McCormick and Bal, 1997) by a 6^th^ degree polynomial to offer a slow initial increase and then a sharp raise mimicking T-current before the burst spikes. After this stage, the burst is approximated by a densely packed spike-like pattern between 0 and 1. For visibility, the burst spike peaks are slightly decreasing sequentially with no effect on the model activity dynamics. We therefore approximated the activity of TRN neurons recorded intracellularly, in vivo, which communicate with each other through GABA_**A**_ receptor-mediated synapses, inducing chloride ion channel-mediated IPSPs. The chloride reversal potential is ∼ -71 mV, which is relatively depolarized compared to the -78 mV resting membrane potential of TRN neurons resulting in IPSP-triggered low threshold spikes (LTSs) (Bazhenov et al., 1999). We approximated the -78 to -71 mV dynamic leading to LTS by a 6^th^ degree polynomial (slow to fast upward curving up before burst in Fig. 2A).

**Figure 2.**
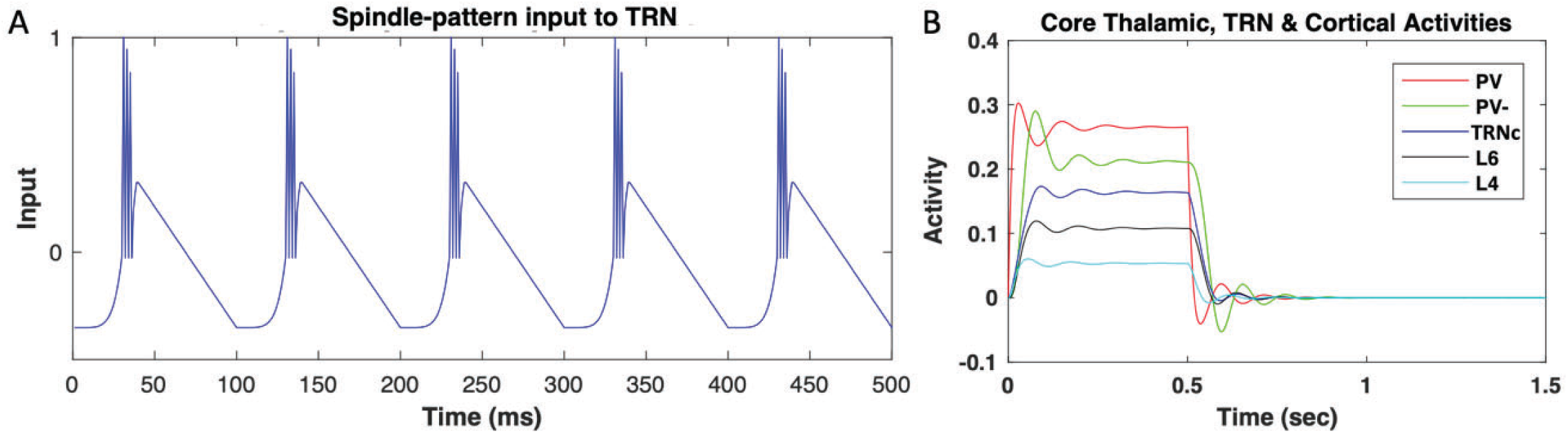
Model inputs and response. **A)** Model TRN input resembling spindle burst temporal pattern. Spindle generation in the model is based on inducing TRN neurons by inputs resembling the spindle spike temporal patterns, which can entrain the model thalamocortical loop depending on the tendency of rebound depolarization and oscillation. Throughout the reported simulations, we chose a 10 Hz (100 ms inter-burst interval) spindle pattern input with a 500 ms duration to induce the model TRN units. To set the TRN spindle inducing pattern as similar to a physiological spindle as possible, we considered Figure 2B of (McCormick and Bal, 1997) and approximated the initial membrane potential raise and then an accelerated upturn (T-current) before the spikes burst by a 6^th^ degree polynomial. We also approximated the bursts by densely packed spike-like pattern between 0 and 1 to have the model neuron activities within the normalized values throughout the simulation. **B)** Model response to 0.5 sec tonic input. Each layer neuron activity is amplitude scaled to prevent overlap for clear visibility. During the input onset, each model neuron depolarizes, reaches its peak, reverses, and after a few oscillations settles. After turning off the square input at 0.5 sec, the activities drop sharply towards baseline followed by hyperpolarization, rebound depolarization, and oscillations to finally settle on the resting state. Response to a 0.5 sec square input illustrates the dynamics of the rate-based model and its tendency for rebound depolarization.

### TRN induced IPSPs in TC neurons and rebound

In the model circuit of Fig. 1, TRN neurons project to the excitatory thalamic neurons in core (PV+) and matrix (CB+) by inhibitory connections (representing GABA synapses). When model TRN neurons receive the spindle inducing input (Fig. 2A), they generate inhibitory postsynaptic potentials in TC neurons (Luthi, 2014; McCormick and Bal, 1997). The postsynaptic potential in TC neurons induces a hyperpolarizing H current leading to cation low-threshold Ca^2+^ channels T-currents before the rebound bursts. The bursts of TC neurons activate the TRN neurons, through the excitatory synapses of the model circuit (Fig. 1, representing glutamatergic input to glutamate receptors of TRN neurons). Figure 2B shows the mutual interaction of model TRN and TC (i.e., PV+) neurons induces rebound activity of TC neurons after hyperpolarization (see for example red PV+ curve right after 0.5 sec). The rebound in the simple compartment rate-based model stems from the decay term -*Av* in equation 1 which brings the hyperpolarized and/or depolarized *v* back to zero with the rate *A* combined with the excitatory and inhibitory neurons’ interactions in the model circuit. Although the simplified mechanisms in the model are by far less sophisticated than the actual physiology of membrane voltage modulation through the variety of hyperpolarization activated cationic H-currents, low threshold Ca^2+^ channels T-current, neurotransmitter gated and voltage gated channels, they nevertheless, approximate the hyperpolarization and rebound of TC neurons.

### Data analysis

Figure 2B illustrates oscillation tendency of the model core thalamocortical loop; after the input is discontinued, activity drops sharply to rest, further reduces toward hyperpolarization, after which activity rebounds toward depolarization. Such a dynamic can cycle a few more times and the more cycles indicate more rebound depolarizations after input shutdown, which are substrates for spindle sustaining. Therefore, we define spindle tendency index (*STI*) as the total duration (in seconds) of the sequence of depolarization rebounds after hyperpolarization. We consider 1% of maximum activity level of a model neuron as the threshold for the peak of a depolarization to be counted as a rebound for inclusion within the hyperpolarization-depolarization duration after input shutdown. We considered the unit of time in seconds because average spindle duration is ∼ 1 second and therefore the number of seconds conveys how many times of an average spindle duration a hyperpolarization-depolarization sequence lasts as an indicator of spindle tendency. For example, in Fig. 2B, the duration of the above threshold hyperpolarization-depolarization sequence after the input shutdown is 0.37 seconds; hence, *STI* = 0.37. The index enables us to compare the tendency of model core, matrix, and mix TC circuits to sustain spindles, as reported in the results section.

## Results

### Rate-based model simulations: ability to sustain and propagate activity

We constructed a computational TC circuit that included core and matrix components with an optional and variable cortical L5 to TRN projection (L5-TRN ON/OFF). Based on the features of TC circuits, our model was able to simulate relay and filtering of signals and could propagate and sustain spindle oscillations. As such, throughout the simulation results reported in the following sections, the state of the network supporting the spindle was based on the presence and amplitude of single *vs* sustained oscillations. To evaluate the readiness of the network to generate or maintain spindles, we gave the model TRN neurons of the model core TRNc the spindle-like input depicted in Figure 2 for 500 ms with starting time at 0 reflected on the x-axis. As indicated in Figure 2 and elaborated in the Data Analysis section, we shut down the spindle-like input at time 0.5 sec. After the spindle input induction, we left the network on its own, to see how the model TC loop, could sustain post spindle oscillations (sequential hyperpolarizations and depolarization rebounds) to estimate the STI. STI is a relative measure of network spindle tendency that can be used to compare the model core, matrix, and mix in supporting spindle generation and sustaining. Using STI, we can also compare model spindle tendency with and without layer 5 corticothalamic projections to TRN, local thalamic inhibition, and with mixing of parallel loops.

### Impact of L5 to TRN_M_ connections

Projections from L5 to TRN have only recently been shown for some cortical areas and at variable levels [e.g., (Prasad et al., 2020; Zikopoulos and Barbas, 2006)], with L6 projections to TRN being considered the norm, and several studies showing absence of direct L5 synaptic terminals in TRN [e.g., (Bourassa and Desche^nes, 1995)]. This prompted us to investigate the effects of direct L5 to TRN_M_ projection on spindle dynamics. Figure 3A shows the spindle dynamics in matrix TC loop when the connection between model L5 and TRN_M_ is absent. We considered this condition the baseline. In Figure 3B-C, we turned on synaptic connections between L5 and TRN_M_ at a level equivalent of 5-10% of L6-to-TRN_**C**_ model connection strengths. Compared to Figure 3A, the single compartment voltage dynamics of matrix TC loop model neurons in 3B-C showed a temporal extension of hyperpolarization/depolarization beyond 0.5 sec (when the spindle like input to TRN was shut down) that gradually increased with increased strength of L5 to TRN_M_ input (from Figure 3B to Figure 3C). This suggests that the presence of direct input from L5 to TRN promotes temporal sustaining of spindles.

**Figure 3.**
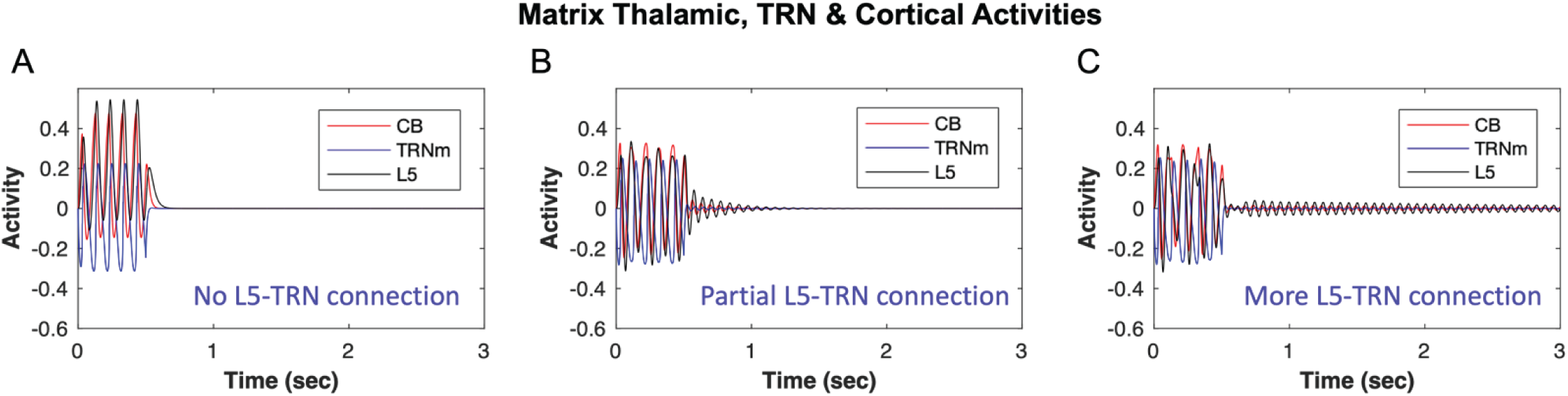
The effect of Model TRN input resembles spindle temporal pattern. The model is based on direct and indirect evidence for the presence of L5 neuron projections to TRN_M_. **A)** No direct L5 to TRN_M_ results in STI < 0.1, which does not promote spindle tendency in the network. **B)** In comparison, implementation of partial, small L5 to TRN_M_ connection (equivalent to 5% of L6 to TRN_**C**_ model connection strength) yields STI = 0.57, which shows a facilitating effect of direct (L5 to TRN_M_) connection in promoting spindle tendency of the network. **C)** the efficacy of L5-TRN connection is further increased from 5% to 10% resulting in STI > 1 that can contribute to the elongation of spindling time and can facilitate continuous or repetitive spindles instead of isolated ones.

### Spindle wave propagation

Kim et al., (Kim et al., 1995) recorded simultaneously from multiple sites in the ferret dorsolateral geniculate nucleus (LGNd) in vitro and found that spindle oscillations propagate across the slice. The hyperpolarization and depolarization waves in Figure 4A show the same trend; spindling tendency propagated across locations where model array neurons numbered from 1-200 (y-axes) are distributed. Figure 4B shows the hyperpolarization-depolarization dynamics of model TC neurons for four locations indicated in Fig. 4A: 1^st^ being center (neuron #100), 2^nd^ location (neuron #113), 3^rd^ location (neuron #128), and 4^th^ location (neuron #147). Following 1^st^ to 4^th^ locations in order shows that the initiation of spindle tendency propagated from the 1^st^ to the 4^th^ location. This provided a mechanistic platform to further investigate the propagation dynamics based on the involvement of different constituent parts of the TC loop. The example in Figure 4B is based on a mixed core-matrix circuit with only core (PV+) neurons’ membrane potentials shown.

**Figure 4.**
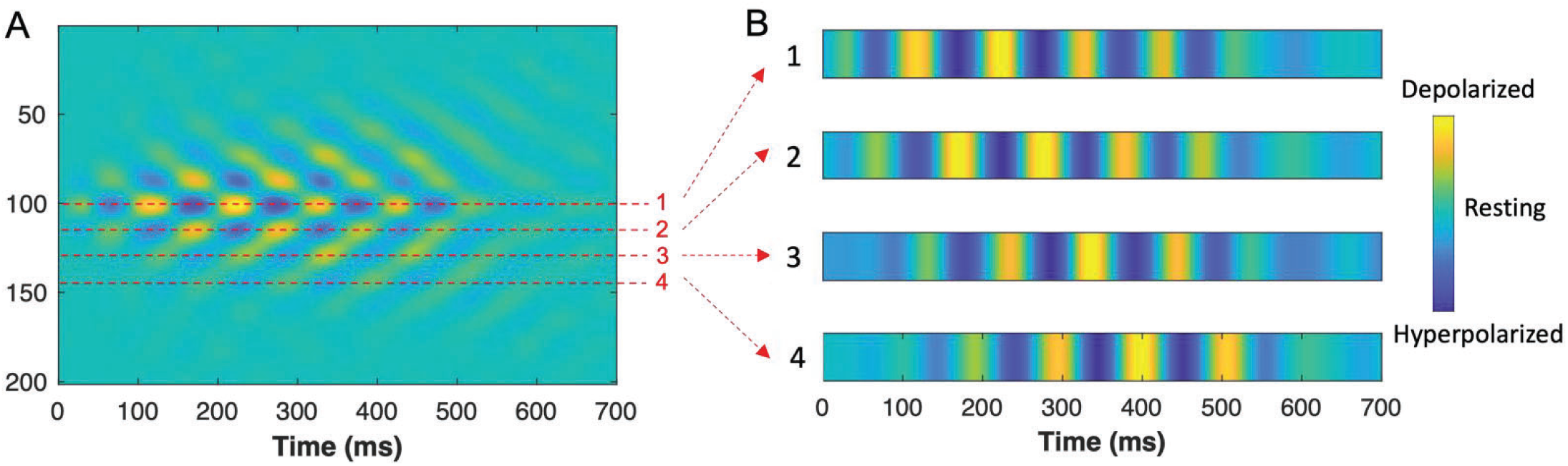
Spindle wave propagation across model neuron arrays. **A)** The activity of 200 model TC neurons (PV+) over time. Yellow, depolarization; green, resting potential; and blue hyperpolarization. Model recording sites are indicated by 1, 2, 3, and 4 corresponding to neurons 100, 113, 128, and 147, respectively. **B)** From top to bottom, 4 recording locations 1-4 from the model neurons are shown. Similar to the (Kim et al., 1995) recording simultaneously from multiple sites in the ferret LGNd, spindle oscillations propagate across locations 1-4 in the model.

### Impact of thalamic local inhibition on spindle dynamics

By tracing the behavior of the model TC neurons under different conditions, we evaluated the tendency of the network to sustain or terminate spindles. Figure 5A shows the model core neuron activities when there is no local inhibition in the thalamus (no PV-→ PV). Figure 5B shows the effects of local inhibition in the thalamus on the oscillatory TC activity. The presence of local inhibition in thalamus indicates a higher spindle probability in the model neurons of core TC loop, as seen in the comparison of the traces of activity of PV and L4 neurons on and after 0.5 sec, which lead to STI >1 in Fig. 5B compared to STI = 0.55 in Fig. 5A. Similarly, local inhibition (CB-) in the matrix TC loop increased the spindle probability when the L5-TRN_M_ connection was present. With no L5-TRN_M_ (STI < 0.1, Fig. 3A), presence or absence of local inhibition did not make a difference and STI remained < 0.1, indicating the necessity of L5-TRN_M_ connection for the gating effect of local inhibition in promoting spindling tendency.

**Figure 5.**
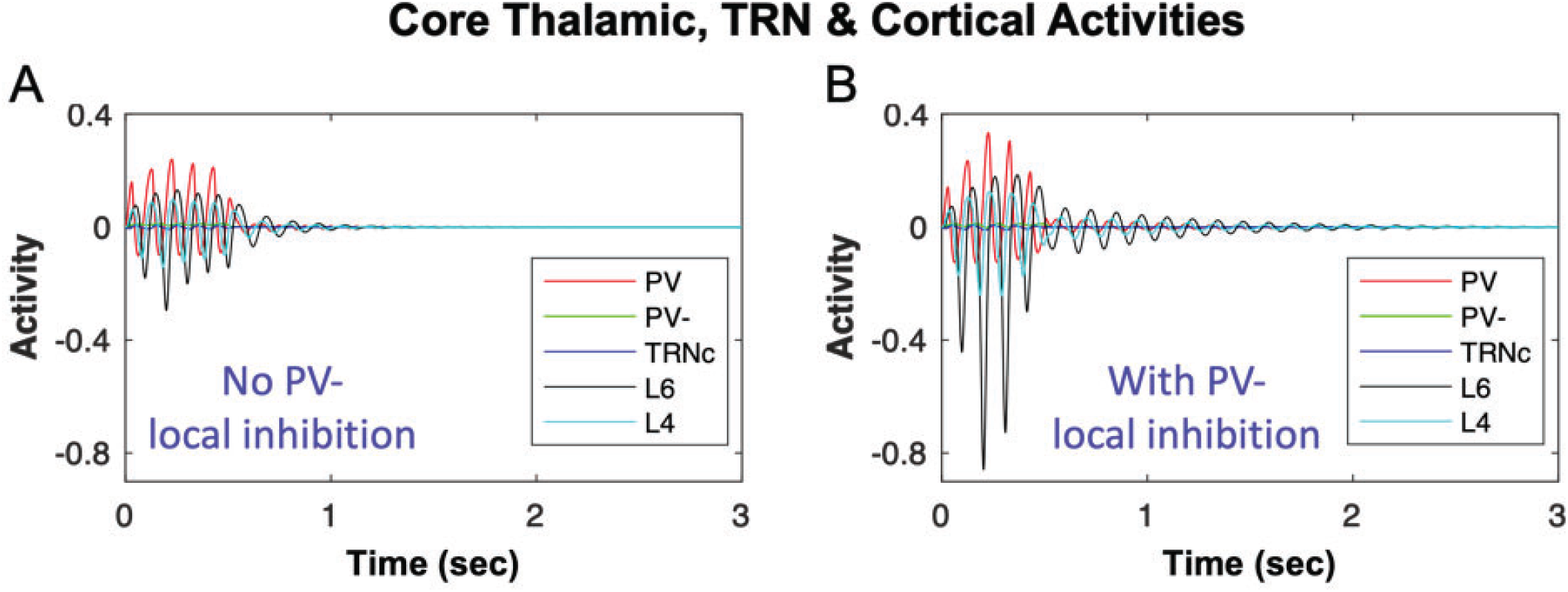
Impact of local thalamic inhibition in biasing the network toward sustaining or terminating spindles. **A)** shows the dynamics of core model neurons with the absence of local inhibition (no PV-inhibition of PV+); and **B)** with the presence of local inhibition. The spindle-like input (shown in Figure 2) is ON for 0.5 sec starting at time 0 and turned OFF at time 0.5 sec. By tracing the behavior of the model TC neurons, we evaluate the tendency of the network to sustain or terminate spindles. Tracing the activity of PV and L4 neurons in (A) and (B)on and after 0.5 sec indicates longer duration and higher amplitude of hyperpolarization and rebound polarization in (B) compared to (A), indicating a higher spindle probability in the model neurons of TC loop with the presence of local inhibition in thalamus.

### Impact of TRN inhibition of thalamus on spindle dynamics

Following the same approach, we tested the effect of different levels of TRN inhibition and cortical feedback on the TC loop tendency to generate and sustain spindles (Fig. 6A-D). Low vs. high levels of TRNc input to model excitatory core PV thalamic neurons (Fig. 6A-C) showed that during the initial spindle-like input to model TRNc (as shown in Figure 2) from time 0-0.5 sec, the amplitude of hyperpolarization and rebound in core model neurons were higher with increased TRNc inhibition of PV neurons. After the spindle-like input to model TRNc was shut down (after 0.5 sec), the sequence of hyperpolarization and rebound events due to model core TC loop sustained longer over time with higher TRNc inhibition of PV neurons (Figure 6C with STI >1) compared to lower TRN inhibition (Figure 6A with STI = 0.55). A similar trend was present for matrix TC loops; higher TRN_M_ input to excitatory thalamic neurons (CB+) increased the STI.

**Figure 6.**
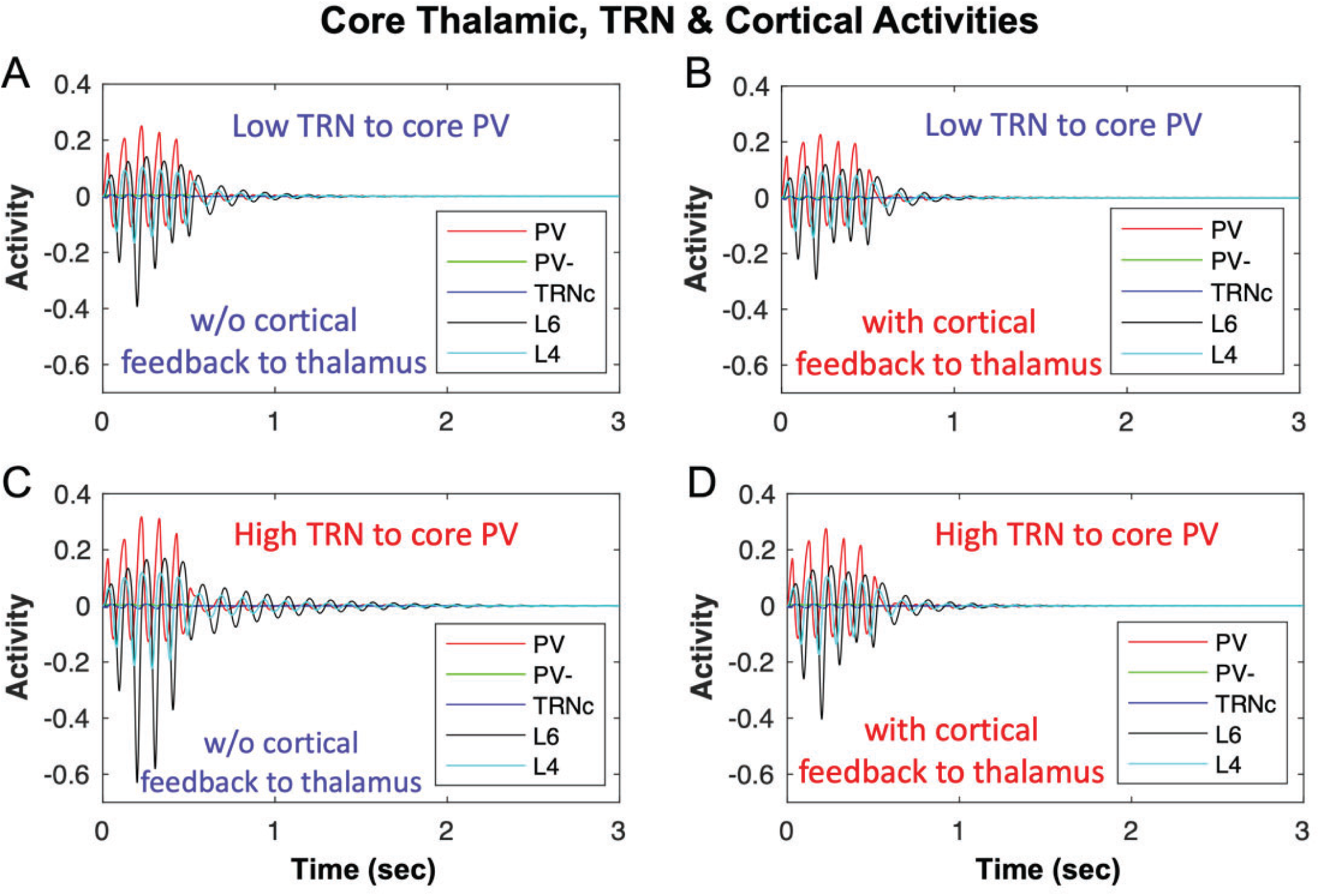
Impact of TRN inhibition and cortical feedback to thalamus in biasing the network toward sustaining or terminating spindles. **A)** shows the dynamics of core model neurons without cortical feedback to thalamus (STI = 0.55) compared to **B)** with cortical feedback to thalamus (STI < 0.55), indicating a decreased spindling tendency. **C)** similar to (A), but with increased TRN inhibition of PV, which increased STI to > 1 compared to (A) with STI = 0.55; **D)** similar to (C) but with cortical feedback to thalamus, which reduced STI to 0.55 from >1 in (C).

### Impact of cortical input to thalamus on spindle dynamics

Corticothalamic feedback in the TC loop controls sleep spindle duration in vivo (Bonjean et al., 2011). In line with this, cortical feedback in our model was a key player in the spindling time and tendency to keep spindles unitary, repetitive, or sustained. In Figs. 6A, C we illustrate an example of the dependence of spindle dynamics on synaptic interactions within the thalamo-reticular loop, without any cortical feedback. Comparison of Figure 6C (STI > 1) with 6D (STI = 0.55) can provide an example of the impact of model cortex feedback to TC neurons. We found that the effect of cortex was toward reducing the spindling time, and after the spindle-like input was shut down at 500 ms, there was less tendency for spindles, in the form of fewer hyperpolarization and rebound depolarization events. Cortical feedback can be channeled in two forms: cortico-thalamic (shown here in Fig. 6, L6 → PV) and cortico-reticular (not shown). Consistent with what (Bonjean et al., 2011) have shown (see their Figure 4), we also found that in our single compartment voltage rate-based model, more feedback from L6 to PV (cortico-thalamic) reduces the spindle duration (Figure 6D; compare with 6C with no cortical feedback). On the other hand, greater cortical feedback to TRN in our model leads to greater input from TRN to thalamus and has the opposite effect, increasing spindle tendency, in line with previous findings (Bonjean et al., 2011). The effects of the model matrix cortical feedback to CB and TRN_M_ were similar to the effects observed in the model core.

### The impact of mixed core and matrix circuits on spindle spatiotemporal dynamics

Most TC loops in primate brains appear to include a mixture of core and matrix components (Jones, 2007; Müller et al., 2020; Piantoni et al., 2016; Zikopoulos and Barbas, 2007a). For this reason, we compared the spatiotemporal dynamics of spindles in the mix circuit with those of isolated core and matrix TC circuits (Figure 7). There are multiple ways of mixing core and matrix circuits, namely, through thalamus, cortex, or both. In thalamic mixing (Figure 7), we connected model PV and CB to TRN_M_ and TRN_C_, respectively. In cortico-thalamic mixing (not shown), we connected model L6 and L5 to TRN_M_ and TRN_C,_ respectively. Finally, we also included in the model laminar interactions occurring at the cortical level, which constitute the most widely inferred, modeled, and studied mixing of core and matrix TC circuits. The mixed spindle spatio-temporal dynamics turned out to be a hybrid of the core and matrix spindles in all cases. These results highlight the impact of circuit connectivity on the core and matrix spindle patterns (Piantoni et al., 2016), which blended seamlessly in mixed designs, despite the constituent regional neural unit differences in core and matrix TC loops.

**Figure 7.**
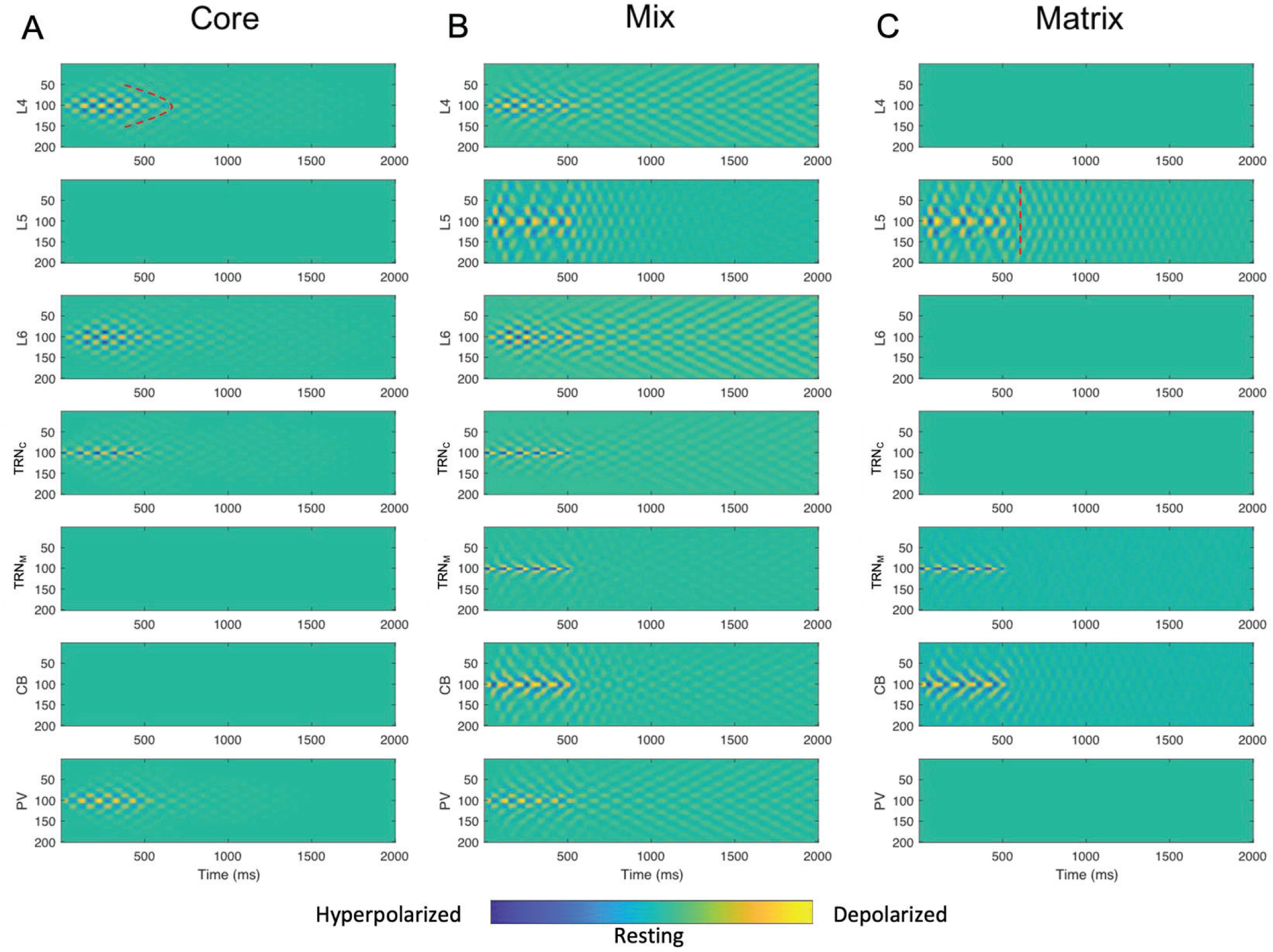
Core, Matrix, and mixed TC model spatiotemporal dynamics. In (A) only model L6, TRN_C_, and PV (core TC loop) were involved in spindle entrainment, induced by inputting spindle like input (Figure 2) to TRN_C_, while L5, TRN_M_, and CB (matrix) remained inactive. In (B), the mix (50-50% thalamo-reticular mixing) of core and matrix, engaged all model areas, and in (C), only model L5, TRN_M_, and CB (matrix TC loop) were involved in spindle entrainment, induced by inputting spindle like input (Figure 2) to TRN_M_, while L6, TRN_C_, and PV (core) remained inactive. Spindle like input (Figure 2), was delivered to neuron # 100 (y-axis) in the TRN_C_ and TRN_M_ array of model neurons, lasting for 500 ms and was then shut down. The presence of colormap ripples in the figure panels are equivalent to sequences of hyperpolarization and depolarization events shown in previous figures and indicate spindle tendency of the network (beyond 500 ms based on the TC model dynamics). Comparison of corresponding panels in (A) and (C) revealed a broader spatial spread and temporal extent in matrix than core. Notably, the mix circuit spatiotemporal dynamics of spindles in each site was a hybrid of core and matrix spindles in terms of broadness of spatial smear and temporal extent. In (A), the model core cortical layer 4, L4, showed a focal tendency for spindle sustaining indicated by a curved dotted marking. Compare with (B) model matrix cortical layer 5 (L5), in which the spindle tendency across model neurons after propagation appeared synchronized and therefore the depolarization iso-amplitude traces aligned vertically at each time bin after spindle propagation for a few hundred ms. A vertical dotted mark as an example is shown in (C) L5.

Figure 8 shows a range of cortical core/matrix mixing ratios from 80% core / 20% matrix to 20% core / 80% matrix in four steps. The level of core and matrix involvement influenced the spatiotemporal dynamics of spindling tendency in the network. Mixed TC loops were mostly influenced by matrix toward a diffuse spatial and smeared over time spindling tendency (hyperpolarizations and rebound depolarizations). Neurons of the model matrix TC loops (e.g., in L5 or CB) influenced the spindling of core neurons (e.g., in L4 or PV) in the mix, while retaining a relatively unaffected diffuse and smeared in time pattern of spindling.

**Figure 8.**
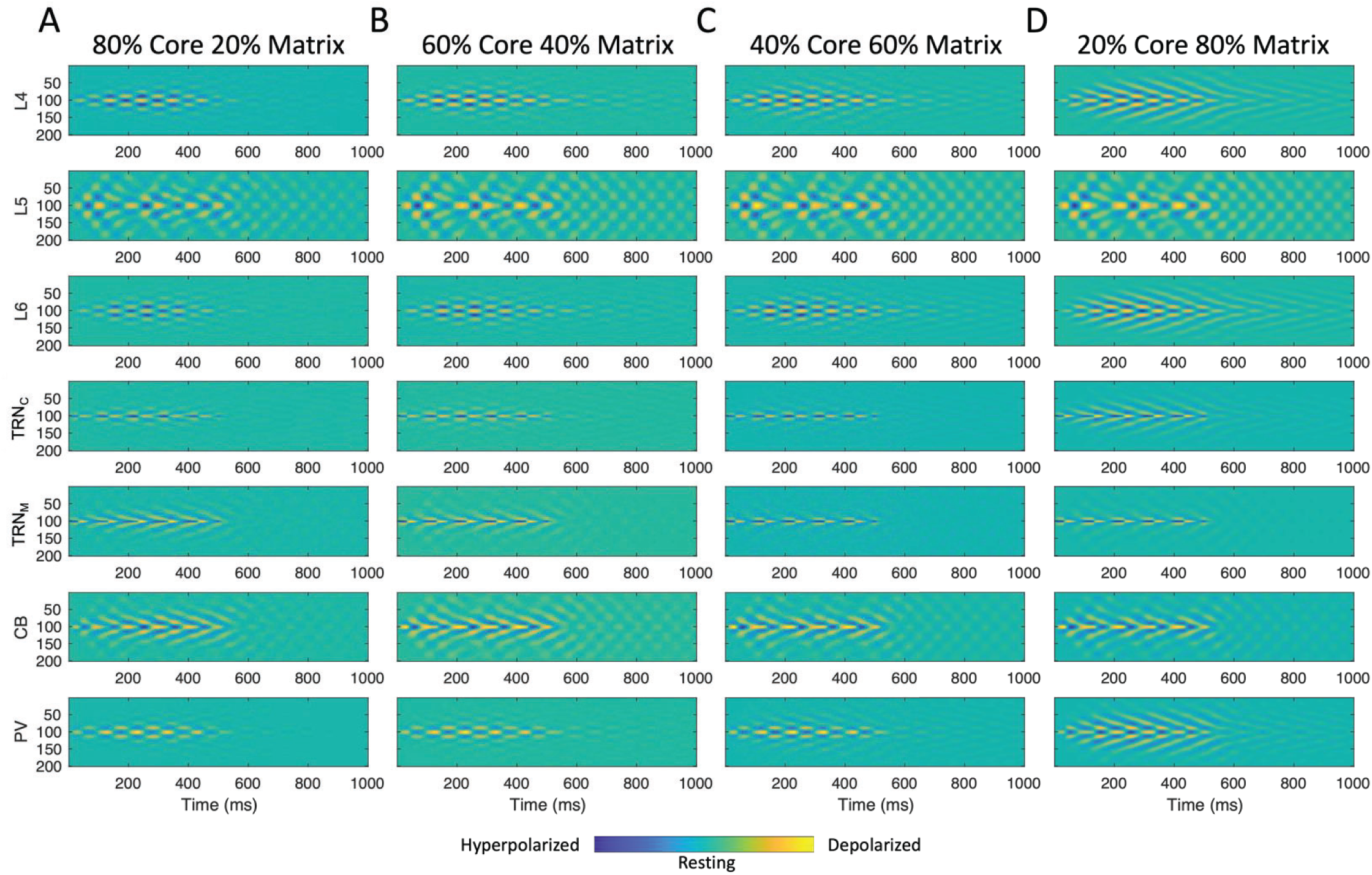
Spatiotemporal dynamics of cortical mixed TC model with different ratios of involved Core and Matrix. **(A-D)** illustrate the spatiotemporal dynamics of TC neuronal activity with different ratios of mixed core/matrix: 80/20, 60/40, 40/60, and 20/80, with mixing occurring at the level of the cortex. The higher the ratio of core/matrix, the more core-like were the spatiotemporal dynamics, i.e., focal in space and less smeared over time. Conversely, the higher the ratio of matrix/core, the more matrix-like were the spatiotemporal dynamics, i.e., more diffuse in space and smeared over time. This trend could be seen in all (in particular core) regions (L4, L6, TRN_C_, and PV) involved in the mix TC loop; the spatio-temporal dynamics of matrix specific regions (L5, TRN_M_, and CB), more or less, remained the same, i.e., diffuse and smeared across different ratios of mixing, as if matrix kept its diffuse spatiotemporal dynamics due to its wider horizontal (within cortical laminae) connectivity and depending on the ratio of core/matrix in the mix, the activity of core regions could be more, or less matrix-like.

Thalamic or cortical mixing produced relatively similar results. The difference was the granularity of the impact of mixing. In cortical mixing, the effect was fine grained (Figs. 8 & 9), in which the graded increment of efficacy or ratio of mixing resulted in graded change in the spatiotemporal dynamic of response. On the other hand, in thalamic mixing, the initial increment of efficacy or ratio of mixing brought the spatiotemporal dynamic of mix to the max/plateau level quickly, acting as a coarse tuning knob (Fig. 7).

## Discussion

We developed a circuit-based TC computational model, with distinct core and matrix loops, based on detailed neuroanatomical organization and connectivity data of TC circuits in primates that can simulate sleep spindles. We additionally simulated mixed TC loops through novel multilevel cortical, thalamo-reticular, and cortico-reticular mixing to investigate the functional consequences of different ratios of core and matrix node connectivity contribution to spindle dynamics. Importantly, our rate-based model, for the first time to our knowledge, included and examined the role of local thalamic inhibitory interneurons, and direct layer 5 projections of variable density to TRN (L5-TRN) and thalamus. Simulations of our rate-based model circuit showed that: a) increased local inhibition in the thalamus or b) increased TRN inhibition of core and matrix thalamic neurons could enhance spindle generation and sustain spindle activity for longer periods; c) the nature of spindles in matrix was more diffuse compared to core, with the mix type showing intermediate properties in agreement with hypotheses that spindles can be classified in core-generated, matrix-generated or mixed types, depending on the neuroanatomy of pathways involved in their generation (Piantoni et al., 2016); d) the L5-TRN projection enhanced spindle generation and propagation; e) spindle power could be modulated based on the level of cortical feedback and involvement in model core vs. matrix; f) matrix TC spindles synchronized their spatial propagation early on, whereas core TC spindles tended to remain spatially focal. In the mix model, the activity of core neurons synchronized and inherited matrix synchronization.

### Connections from cortical L5 to TRN are necessary for heterogeneity of spindle oscillations, especially for matrix and mix spindle generation

Projections from L5 to TRN have only recently been shown directly or indirectly for some cortical areas and at variable levels (Hádinger et al., 2022; Prasad et al., 2020; Zikopoulos and Barbas, 2007b, 2006). In primates, several thalamic nuclei, especially those connected with prefrontal cortices, receive anywhere from 20-50% of their cortical projections from layer 5 pyramidal neurons (Xiao et al., 2009). These thalamic nuclei, which receive projections from pyramidal neurons in cortical layers 5 and 6, participate in matrix TC loops, in addition to typical core TC circuits (Zikopoulos and Barbas, 2007a). These robust projections from layer 5 terminate as axonal branches with a mix of small and large axon terminals that form synapses with thalamic projection neurons and interact with ionotropic glutamate receptors (Reichova and Sherman, 2004; Zikopoulos and Barbas, 2007a). In primates, these types of axons that contain large or a mix of large and small axon terminals are also found in TRN regions that are connected with prefrontal cortices, account for about 38% of all prefrontal axons, and most likely originate in cortical layer 5. These types of axon terminals are not seen in pathways from sensory cortices to TRN, in primates (Zikopoulos and Barbas, 2006).

Our simulations showed that the cortical projection to TRN is a critical circuit component that determines the spatiotemporal dynamics of spindle propagation and underlies the heterogeneity of spindles, observed in functional and physiological studies of the human brain (Bonjean et al., 2012, 2011; Mak-McCully et al., 2017, 2014). The L5-TRN connection, in particular, appears essential for the generation of matrix and consequently mix spindles. With regard to mix spindle dynamics, we examined, for the first time, mixing of the core and matrix connections at multiple levels. Our findings suggest that mixing, both at the thalamo-reticular level (CB+ and PV+ thalamic projection neurons sending their signals to TRNc and TRNm, respectively), or at the cortico-reticular level (L5 and L6 pyramidal projection neurons innervating TRNc and TRNm, respectively), can act as a coarse knob for spindle control. That is because the spindle generator (TRN) is targeted in the mix, and, as such, the spindle dynamics change significantly, even after low levels of mixing; hence, we have a narrow dynamical range for mixing. Conversely, cortical mixing can act as a fine knob. This is because, mixing at a level not directly involved in the generation of spindles (i.e., cortex), can be done with a broad dynamical range (fine knob), because the cortex is entrained by the spindle generator level (thalamo-reticular), rather than being a generator itself. Cortical mixing is likely the prevalent mode of mixing of TC circuits, because it relies on interlaminar connectivity that constitutes the main component of cortical microcircuitry, i.e., projections from neurons in L3/4 to pyramidal neurons in L5, or core cortical nodes in L4 and L6 receiving L1-3 signals that are also influenced by a wide kernel of matrix inputs (Krishnan et al., 2018; Thomson, 2007; Yoshimura et al., 2005). To this effect, recent physiological studies have highlighted the laminar heterogeneity of human sleep spindles (Hagler et al., 2018; Krishnan et al., 2018; Thomson, 2007; Yoshimura et al., 2005), but also the extensive mixing of spindles across layers of the human cortex (Ujma et al., 2021).

### Local thalamic inhibition increases ability to generate, sustain, and propagate spindle activity

Our simulations showed that increased local inhibition in thalamic nuclei, which can hyperpolarize either core or matrix TC projection neurons, could enhance spindle generation and sustain spindle activity for longer periods. Importantly, GABAergic local circuit neurons are abundant in all the dorsal thalamic nuclei of primates, and may constitute about 40% of all neurons in the human thalamus (Arcelli et al., 1997; Hunt et al., 1991; Smith et al., 1987), but are not present in all the thalamic nuclei of different mammalian species, and are virtually absent in the thalamus of rodents, with the exception of few first-order thalamic nuclei like the lateral geniculate nucleus (Arcelli et al., 1997; Barbaresi et al., 1986; Gabbott et al., 1986) and the ventral posterior nuclei (Simko and Markram, 2021). The importance of local thalamic inhibition has not been explored in most experimental and computational studies of TC circuits or sleep spindles, with few exceptions (Arcelli et al., 1997; Joyce et al., 2022; Sherman, 2004; Timbie et al., 2020). There is agreement that local inhibition, which is especially abundant in primates, adds complexity to the synaptology and circuitry of the thalamus, in part by facilitating widespread formation of triadic synaptic glomeruli in most thalamic nuclei (Arcelli et al., 1997; Joyce et al., 2022; Sherman, 2004; Timbie et al., 2020), and likely affects the functional properties of TC circuits that can modulate attention, memory consolidation, and spindle tendency.

Importantly, the ubiquitous presence of GABAergic local circuit neurons in the primate thalamus may provide alternative wiring avenues for the construction of open loop circuitry linking TRN with the thalamus (Fig. 1B; also see for example Fig. 9 of (Barbas and Zikopoulos, 2007) and Figs. 5 and 9 of (Zikopoulos and Barbas, 2007b). Several studies suggest that the open loop model of TRN-thalamic interactions, in which TRN neurons are excited by a thalamic neuron and, in turn, inhibit a different thalamic neuron or an inhibitory interneuron, can create a tunable filter that may be modified by extra-TRN influences, like cortico-reticular pathways (Brown et al., 2020; Willis et al., 2015). On the other hand, closed loop TRN-thalamic connectivity, in which TRN-thalamic neuron pairs are reciprocally connected, allow a short window for initial excitation of a TC neuron followed by inhibition, preventing summation and faithfully transmitting high-frequency signals to the cortex (Steriade and Deschenes, 1984). Since both circuits are found in the primate thalamus, we modeled TRN-thalamic interactions using a novel Hybrid Loop Design (Fig. 1C) that can flexibly simulate either open or closed loops. The Hybrid Loop Design we implemented represents the spread of TRN-Thalamic connectivity strength using a bilateral Gaussian curve system. Our architecture resembles a closed-loop functionally, when the bilateral Gaussian spread of connectivity strength is spatially symmetric, which means that the resulting Gaussian peak of connectivity strength is to the thalamic neuron which directly excites the TRN neuron. In other words, a closed loop in our model is composed of two open loops in opposite directions (balanced). Conversely, when the connectivity spread is either spatially asymmetrical or follows a shifted Gaussian, then our circuit architecture resembles an open-loop. The Hybrid Loop TRN-Thalamic connectivity design used in this study can be a powerful tool to further explore open vs closed loop architecture and interactions in the thalamus and other systems in future studies.

**Figure 9.**
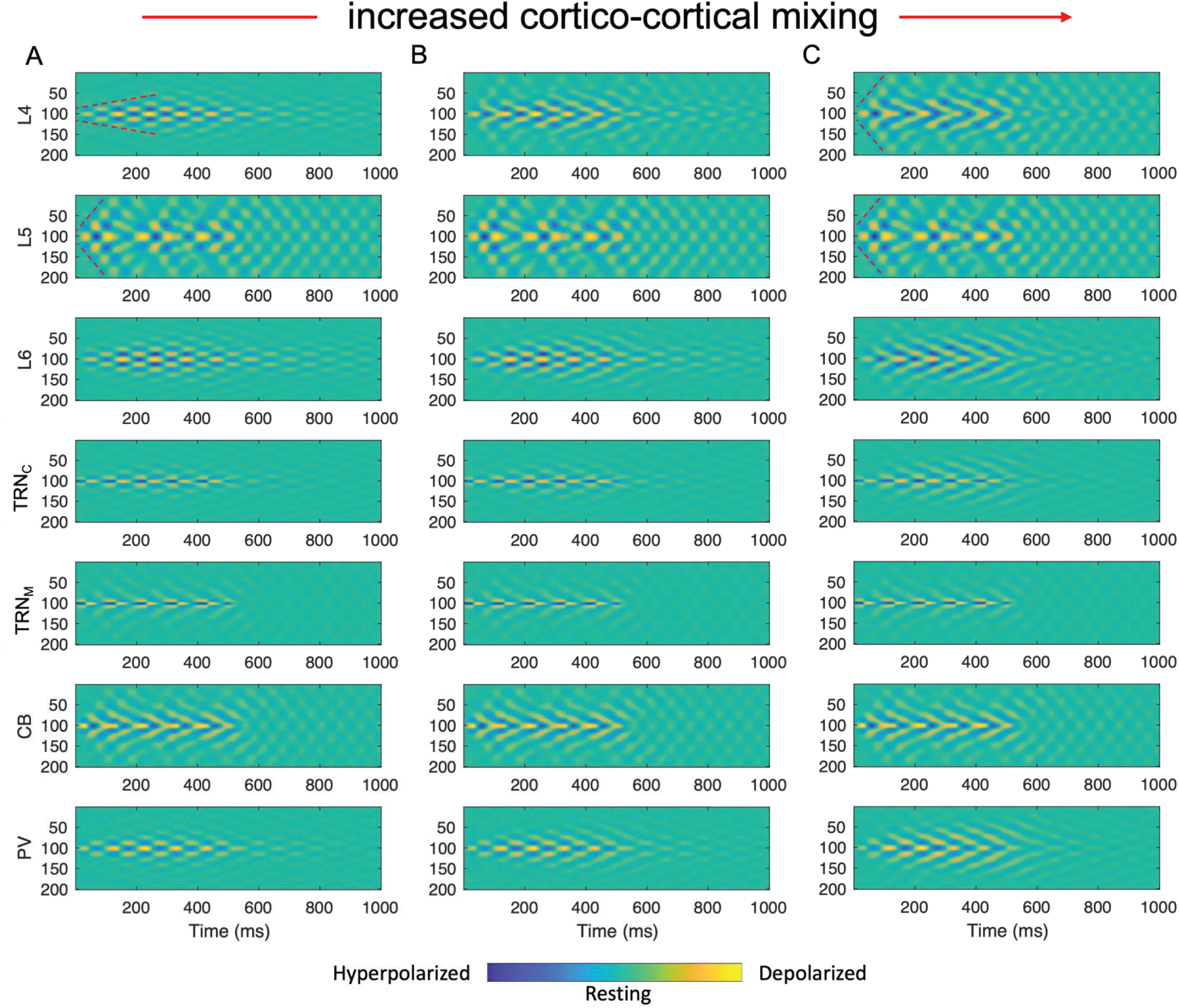
Spatiotemporal dynamics of TC model with different synaptic efficacy of cortical mixing of Core and Matrix. The spatiotemporal dynamics of the TC model plateaued after reaching increased synaptic efficacy in cortical mixing. In **A)** 30%, **B)** 60%, and in **C)** 90% of the plateau synaptic efficacy being used. Similar to Fig. 8, matrix model neurons kept their spatio-temporal dynamics, and the response of core model neurons resembled the response of matrix model neurons, as the efficacy of mixing gradually increased. Model core cortical layer 4 (L4), in 90% mixing level (C), compared to 30% mixing level (A), showed faster activity propagation across neurons indicated by higher slope (i.e., number of neurons per ms) of red dotted marks. This shows that mixing core with matrix impacts core signal propagation rate. On the other hand, matrix fast propagation rate remained relatively the same across different mixing levels of core and matrix, indicated by the same slope of red dotted marks in (A) and (C) L5 panels.

### Increased cortical influence on TRN vs thalamus affects the tendency to generate spindles in opposite ways

Previous studies have shown that TRN neurons are tuned to be more responsive to cortical than thalamic inputs, based on specialized synaptic-receptor interactions (Golshani et al., 2001; Liu and Jones, 1999). This suggests that changes in cortical input may affect differentially TC vs TRN neurons. Our simulations show that increased ratio of core or matrix TC neuron activation, due to increased cortical feedback to TC that is not accompanied by equivalent increase of cortical feedback to TRN, can reduce a reverberating tendency to promote spindle duration and sustaining. In contrast, increased cortical feedback to TRN neurons can reverse the balance, and increase the spindle tendency (not shown), consistent with findings showing that increasing the cortico-reticular conductance (i.e. g(PYR→TRN)) can result in total spindle duration increase (Bonjean et al., 2011), and in line with the spindle generator properties of TRN neurons. However, recent studies have also suggested that depending on the state of cortical and thalamic neuronal activity, high levels of cortical input to TRN may, in some cases, disrupt thalamic spindling (Bonjean et al., 2012, 2011; Mak-McCully et al., 2017, 2014), highlighting the complexity of the TC circuit interactions in primates.

### Implications for brain disorders

Thalamocortical circuits are organized into diffuse matrix and spatially selective core components that are mixed in different ratios throughout the brain (Piantoni et al., 2016). TC circuits can indirectly propagate signals from one cortical area to another, in concert with or in addition to direct corticocortical pathways, forming a framework for rhythmic brain activity, including signal and spindle propagation across cortex (Brown et al., 2020; Halassa and Sherman, 2019; Jones, 2001; Sherman and Guillery, 2013; Sherman, 2012). Moreover, thalamocortical and cortical connectivity influence sleep spindle properties (Krishnan et al., 2018). Therefore, changes in spindle dynamics can be considered a litmus test for typical thalamocortical circuit connectivity and for disruptions underlying neuropathology, because the latter can also impact spindling tendency. For example, medicated individuals with chronic schizophrenia show deficits in spindle density (Ferrarelli et al., 2007; Manoach et al., 2010), which parallel impaired memory consolidation during stage 2 sleep (Göder et al., 2015; Wamsley et al., 2012), and increased thalamocortical functional connectivity (Baran et al., 2019; Manoach and Stickgold, 2019). Moreover, spindle density and duration are significantly decreased in autism spectrum disorders (ASD) and show negative correlation with sleep-dependent memory consolidation (Farmer et al., 2018; Mylonas et al., 2022). Both disorders differentially affect sensory processing, attention, and emotional processing likely involving at variable degrees first-order, primary networks that participate in core TC circuits, or high-order, association networks that participate in matrix or mix TC circuits. Therefore, changes in spindle dynamics can also point to the imbalance of core and matrix involvement in disorders such as schizophrenia and ASD, pinpointing specific TC loops, circuit nodes, cell types, and underlying mechanisms that are likely disrupted. These disruptions are manifested through distinct symptomatology, for example, distractibility and difficulty focusing attention is a hallmark of schizophrenia (Braff, 1993; Luck and Gold, 2008) whereas, difficulty switching attentional focus and sensory over-responsivity are often seen in ASD (Allen, 2001; Cheung and Lau, 2020; Fan et al., 2012; Green and Ben-Sasson, 2010; Keehn et al., 2013; Marco et al., 2011). Future studies can use the modeling framework we developed to further investigate how key circuit organization features in primates including thalamic inhibition, locally or through TRN, and extensive L5-TRN projections, can change spindle generation, and their spatiotemporal propagation. This framework can additionally provide useful insights on sets of neurobiological and connectivity metrics, including genetic and epigenetic modifications of thalamic, TRN, and cortical pyramidal neurons that can be used to infer potential underlying mechanisms of changes in spindle dynamics in disorders such as ASD and schizophrenia.

